# Mid-superior temporal sulcus encodes spatial context and behavioral state in freely moving macaques

**DOI:** 10.64898/2026.04.16.719074

**Authors:** Felipe Parodi, Alessandro P. Lamacchia, Yufan Ye, Pooya Laamerad, Yiting Chen, Kristin L. Gardiner, Sebastien Tremblay, Konrad P. Kording, Michael L. Platt

## Abstract

Primate neuroscience has traditionally studied the brain under highly constrained conditions, limiting our understanding of neural function during real-world behavior. The mid-superior temporal sulcus (mSTS) is implicated in social perception, but its role during unconstrained behavior has not been tested. Here we performed wireless depth-electrode recordings from both banks of mSTS in macaques freely exploring a large three-dimensional arena, combined with 3D pose tracking and behavioral segmentation. Neural encoding models revealed mSTS firing rates were jointly modulated by spatial position, body kinematics, and geometric visual proxies, preferentially encoded in allocentric coordinates but with a shift toward body-centric encoding during vertical exploration. Neural populations carried decodable information about discrete behavioral syllables, with broad temporal generalization and neural similarity that tracked the sequential structure of behavior. Population manifold analysis revealed that the same behavior occupied different regions of population space at different spatial locations, and population dynamics showed structured organization around behavioral transitions. Together, these results suggest that mSTS populations carry joint information about spatial context and behavioral state during natural behavior.

## Introduction

Understanding how the primate brain operates during natural behavior is a fundamental challenge for neuroscience. Classic paradigms – motor restraint, controlled stimulus presentation, trial-based task structures – have yielded deep insights but may systematically obscure neural computations that emerge only during unconstrained action^1,2^. The emerging field of primate neuroethology^3^, enabled by advances in wireless neural recording and computational ethology, now makes it possible to study the brain during the rich, self-generated behaviors that define an animal’s natural repertoire^4–13^. This approach is especially compelling for brain regions whose putative roles were established under constrained conditions. Encoding properties characterized under restraint may capture only the surface layer of neural representations that emerge when animals engage the world freely.

The superior temporal sulcus (STS) is an ideal test case for how constrained paradigms may have narrowed our understanding of what a brain region encodes. Across decades of restrained recordings, mid-STS (mSTS) has been characterized as a visual processing hub for biological motion, action observation, and social perception^14–17^. Body-selective patches in STS are hypothesized to progressively transform body and action representations along a hierarchy from low-level motion features to higher-order social functions^18–20^, including intention attribution and gaze monitoring^21–23^. Recent work has further shown that mSTS is modulated by social prediction^24^ and by partner-action prediction errors with a preference for live agents^25^, suggesting that the region’s function extends beyond passive social perception. Yet this characterization derives from paradigms presenting controlled stimuli to restrained observers. Classic findings that neurons in anterior dorsal-STS do not respond to the sight of the animal’s own limb movements reinforced this observer-centric view^26^. This consensus emerged from paradigms that, by design, constrained the sensorimotor correlations that could be written into the observable record of neural activity.

We hypothesize that mSTS carries richer signals during free movement. This possibility is made plausible by (i) its anatomical position at the convergence of dorsal and ventral visual streams, (ii) its inputs from motion-sensitive areas including MT/MST^27^, and (iii) broader work demonstrating that releasing constraints reveals richer encoding. Recordings in monkeys free to move both their heads and eyes revealed gaze displacement signals in superior colliculus invisible under fixation^28^, while studies of head-restrained animals free to move their limbs showed that movement-associated modulation of primate visual cortex primarily reflects visual reafference^29^ – sensory signals generated by the animal’s own movement. Recent prefrontal recordings in monkeys free to move both their heads and eyes combined with movement quantification demonstrated robust cognitive signals despite substantial movement-related variance^30^. Fully unconstrained wireless recordings have further uncovered even richer cortical representations during individual^8,9,12^ and social behavior^6,11^ (see [3] for review). mSTS has thus far eluded this approach because sulcal cortex is inaccessible to chronic surface arrays, making the recordings technically challenging. Dissociating visual from kinematic contributions also requires high-quality 3D movement reconstruction^31,32^ or head-mounted cameras^33,34^. mSTS is therefore a prime candidate for richer encoding during free behavior, but also one of the most technically demanding regions in which to test it.

Recent theoretical work has argued that brain regions associated with social cognition may be recruited during interactions with others because these areas perform computations that are useful in many settings, not only social ones^35^. If that view is correct, then some of the computations performed by mSTS during social perception should also be detectable during solitary behavior, where the animal must continuously monitor its own position, motion, and changing state in a dynamic environment. We tested this by recording from mSTS in macaques freely exploring a large arena alone.

Here, we report findings from wireless, depth-electrode recordings from both banks of mSTS as macaques freely explored a large open-field arena, combined with 30-camera markerless 3D pose tracking and unsupervised behavioral segmentation. Variance decomposition encoding models revealed that static 3D position in the arena explained more unique variance in mSTS firing rates than any single measured visual or kinematic feature group. In separate reference-frame models based on displacement and velocity variables, encoding was stronger in allocentric (world-fixed) than body- or head-centric coordinates overall, with a shift toward body-centric weighting during vertical exploration. At the population level, mSTS encoded discrete behavioral syllables – brief, recurring actions – in a context-dependent neural population space, and neural dynamics showed structured temporal organization around behavioral transitions. Together, these findings support the view that mSTS activity reflects a joint representation of spatial context and behavioral state during natural behavior, expanding beyond observer-centric models and advancing our understanding of how primate cortex operates during real-world behavior.

## Results

### Freely moving macaques generate rich behaviors suitable for quantifying visual and kinematic signals

The computational role of mSTS during natural behavior remains untested, in part because recording from deep sulcal cortex in freely moving primates presents technical challenges. We developed a wireless recording system enabling untethered neurophysiological monitoring while macaques freely explored a large open-field arena that elicits behaviors inaccessible in restrained paradigms (Figure 1A). Depth electrodes targeting both banks of mSTS, coupled to skull-mounted wireless data loggers, yielded sustained recordings from 1,404 single and multi-units (860 SUA, 544 MUA; Extended Data Fig. 1) across five sessions in two subjects during spontaneous behavior, capturing neural dynamics across the sulcal anatomy for the first time without movement constraints. The hexagonal arena (2.4 m tall, 1.5 m per side) contained scattered foraging materials (seeds, nuts) and multi-level structures that afforded climbing, perching, and overhead suspension. This design enabled testing of whether behavioral variables previously inaccessible in restrained paradigms contribute to neural activity in mSTS.

**Figure 1.**
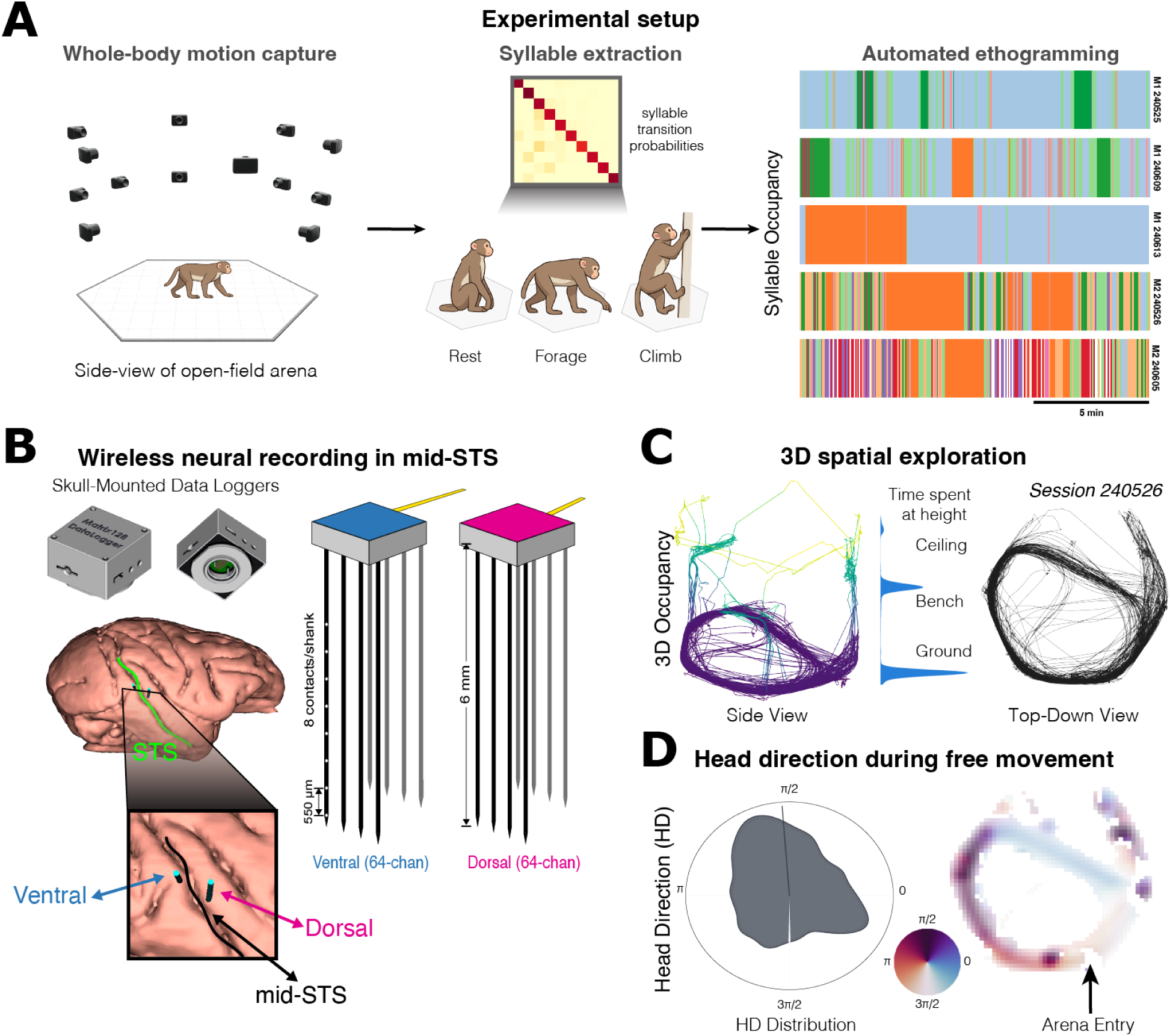
Freely moving macaques generate quantifiable behaviors suitable for dissociating visual and kinematic signals. **a.** Multi-camera motion capture in a large open-field arena (hexagonal, 2.4 m tall, 1.5 m per side) yields whole-body 3D pose estimates from which discrete behavioral syllables are extracted via unsupervised segmentation. Syllable transition probabilities (top) summarize the sequential structure of behavior; example syllable icons (rest, forage, climb) illustrate representative syllables. Right: automated ethograms for five sessions (top three, M1; bottom two, M2; scale bar, 5 min) show time-resolved syllable occupancy. M1 sessions are dominated by rest with limited behavioral diversity, whereas M2 sessions exhibit frequent, rapid transitions across a broader repertoire. **b.** Wireless neural recordings from mid-superior temporal sulcus (mSTS). Left: head-mounted data loggers and chronically implanted depth arrays. Inset: electrode targeting across the ventral and dorsal banks of mSTS. Right: schematic of dual 64-channel depth electrode geometry spanning the sulcus. **c.** The 3D arena elicits rich kinematic variability. Left: center-of-mass trajectory through the enclosure for an example session, colored by vertical position. Center: kernel density estimate of time spent at each height level (ground, bench, ceiling). Right: top-down view of the same trajectory. **d.** Viewing angle during free movement. Left: polar histogram of head-direction distribution. Right: arena trajectory colored by head direction; arena entry indicated.

Quantifying the relationship between behavior and neural activity requires precise, continuous tracking of body and head kinematics. Our 30-camera motion capture system tracked 17 anatomical landmarks at subpixel precision (median reprojection error: 0.35 px), enabling head orientation estimates independent of body position (cf. PrimateFace^36^). Animals exhibited rich behavioral repertoires (Figure 1A), with head velocities ranging from stationary fixations to rapid movements, viewing angles spanning the full azimuthal range, and occupancy across multiple vertical levels, including foraging material on the floor, three benches at the mid-level (approximately 1.5 m above the ground), and a grated ceiling that permitted overhead suspension (Figure 1C and Extended Data Fig. 1). Together, these measurements provide a rich, high-resolution behavioral description suitable for identifying neural correlates of self-motion and visual experience.

To test whether mSTS activity reflects behavioral state beyond visual and kinematic variables, we also characterized discrete behavioral structure using unsupervised behavior segmentation adapted from computational ethology^37–39^. We extracted pose dynamics using 3D pose estimation, and then identified discrete behavioral syllables through dimensionality reduction and clustering. This yielded 12 syllables (median bout duration 1.50 s, IQR 0.43–3.40 s; see Extended Data Fig. 4), from stationary scanning to directed locomotion to overhead exploration. Combined with continuous kinematics, these provide a comprehensive set of behavioral predictors for neural encoding analyses.

### mSTS neurons encode 3D spatial position and self-motion during free behavior

To determine which, if any, behavioral variables mSTS neurons encode during unconstrained movement, we fit ridge regression encoding models to single and multi-unit activity using four feature groups: visual proxies (VIS; gaze direction, distance-to-wall along gaze, and a looming proxy), body kinematics (BODY_KIN; forward/lateral velocity, speed), head kinematics (HEAD_KIN; angular velocities, head-body offset), and 3D spatial position (POS; center-of-mass coordinates in the arena frame) (Figure 2A). Importantly, the VIS features are geometric proxies rather than a reconstruction of the full visual input. They capture line-of-sight geometry to salient arena regions, but not image content (e.g., texture/identity), optic flow, or retinal coordinates. VIS, BODY_KIN, and HEAD_KIN features included temporal lags (±1.0 s) to capture anticipatory and delayed responses; POS entered as a static, unlagged regressor. The full model achieved cross-validated R^2^ > 0.01 in 70.2% of units (986 of 1,404), with encoding particularly robust in M2 (96.6%; Figure 2B) and more variable in M1 (30.7%), likely reflecting M1’s restricted spatial coverage of the arena (Extended Data Fig. 2). These results indicate mSTS activity is broadly behavior-modulated during free movement, motivating finer-grained analyses of individual feature contributions.

**Figure 2.**
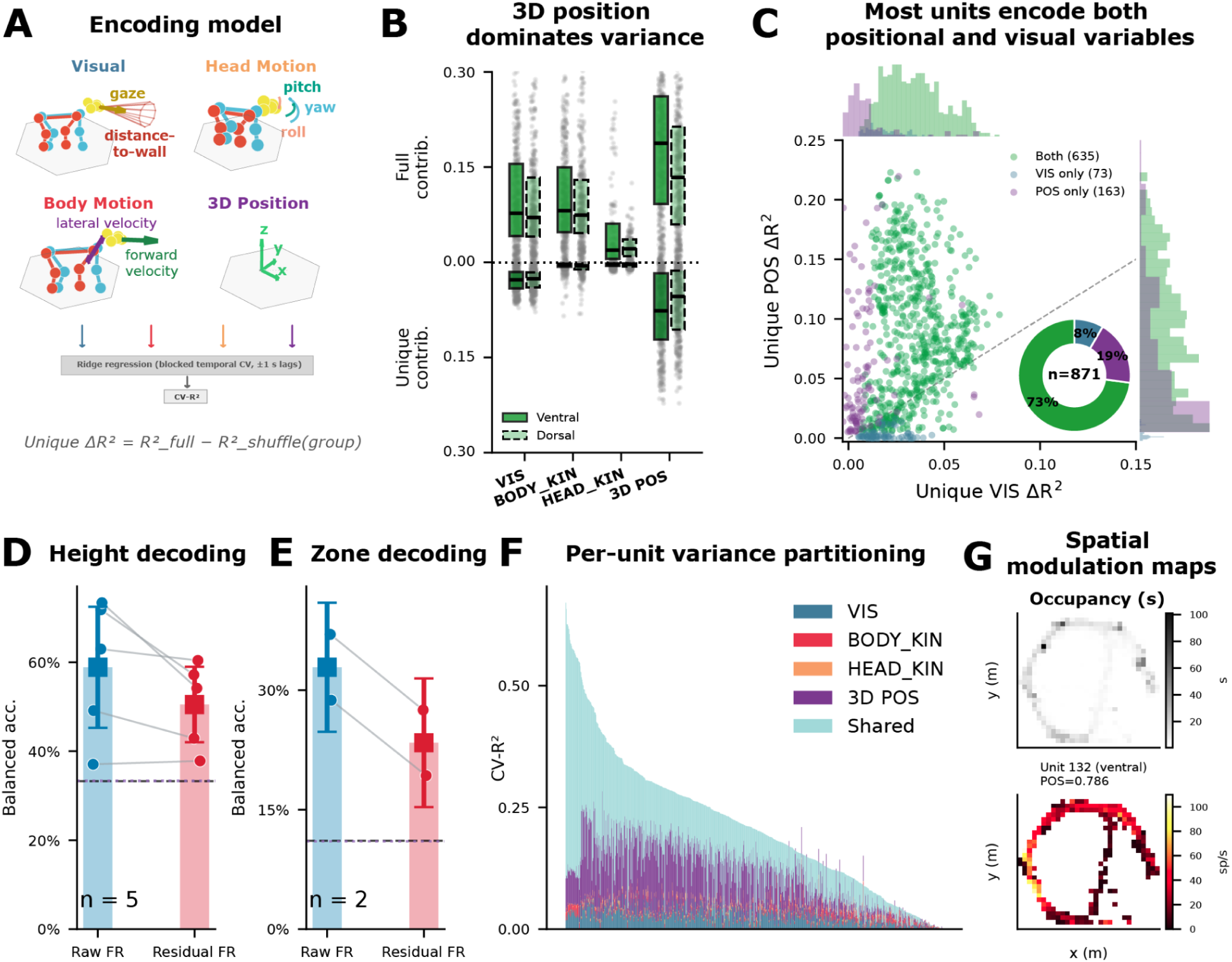
mSTS neurons encode 3D spatial position and self-motion during free behavior. **a.** Schematic of the encoding model. Neural firing rates (500 ms bins) were modeled as a linear function of four behavioral feature groups – visual proxies (VIS), body kinematics (BODY), head kinematics (HEAD), and 3D position (POS) – using ridge regression with blocked temporal cross-validation. Unique variance was quantified by block-shuffling each group and measuring the decrease in cross-validated R^2^. **b.** Full contribution (top) and unique variance (bottom) for each feature group (example decomposition from M2 (n = 843 units, 2 sessions). Boxes show median and interquartile range; individual units shown as gray dots. 3D position in the arena (POS) explained the most unique variance, followed by visual proxy features (e.g., projected gaze; VIS). **c.** Population scatter of unique POS ΔR^2^ (y-axis) versus unique VIS ΔR^2^ (x-axis) for each encoding-positive unit across all sessions (n = 871 of 1,404 total; 2 subjects, 5 sessions). Units colored by significance category: green = significant for both VIS and POS (73%), purple = POS only (19%), blue = VIS only (8%). Marginal histograms show the distribution along each axis; donut inset shows population composition. Most units fall above diagonal, indicating POS-dominant encoding, but the majority carry both signals simultaneously. **d.** Height decoding (3 classes: floor, bench, ceiling; n = 5 sessions). Mean balanced accuracy is shown for raw firing rates and firing rates after regressing out visual, kinematic, and time-drift covariates (bars), with squares indicating mean ± 95% CI; per-session values are overlaid as paired points connected by lines. Decoding is above chance (33%) in both conditions, with higher performance for raw activity (∼59%) than residualized activity (∼51%). **e.** Position decoding (9 classes: height tiers × angular zone; n = 2 sessions, M2 only). Plot conventions are identical to panel D, with an independent y-axis. Decoding exceeds chance (11%) in both conditions, and is higher for raw activity (∼33%) than residualized activity (∼23%). Chance and permutation-chance reference lines are shown in each panel. **f.** Variance partitioning. Stacked bars show the relative unique contribution of each feature group to total explained variance per unit. **g.** Spatial firing maps. Top: top-down view of arena occupancy for session 240526. Bottom: spatial firing rate maps for a top POS-encoding unit (ranked by unique ΔR^2^). Color indicates mean firing rate as a function of 2D arena position.

To estimate how much each feature group contributed beyond the others, we used a shuffle-based analysis that quantified the variance uniquely explained by each group (cf. [^6,30,40^]). 3D spatial position explained the most unique variance (median ΔR^2^ = 0.063), exceeding visual features (0.026), body kinematics (0.004), and head kinematics (0.003) (Figure 2B,F). All feature-group comparisons remained significant after Benjamini-Hochberg FDR correction and were preserved in control analyses (see Methods and Extended Data Fig. 2). These unique variance fractions are modest in absolute terms, as expected in naturalistic paradigms where correlated, high-dimensional regressors distribute shared structure across groups (cf. [^30^]). These results show that mSTS firing rates are more strongly modulated by static 3D position in the arena than by any single measured visual or kinematic feature group.

To assess whether visual and spatial signals are carried by distinct subpopulations, we classified each unit by whether it carried significant unique variance for visual features, spatial position, or both (FDR-corrected, p < 0.05, ΔR² ≥ 0.005). Of the 986 units with R² > 0.01, 871 (62%) carried significant unique variance for at least one group (Figure 2C). Most of these (73%) encoded both visual and spatial signals, with smaller fractions encoding only position (19%) or only visual features (8%). On the per-unit scatter of unique POS versus VIS ΔR², most units fell above the diagonal – indicating stronger spatial than visual encoding – but nearly all carried appreciable variance from both feature groups. Visual and spatial information are thus multiplexed within individual mSTS neurons rather than segregated into distinct subpopulations.

We next asked whether the population carries sufficient spatial information to decode location. We discretized the arena into three height tiers (floor, bench, and ceiling) and decoded tier identity from population firing rates (Figure 2D). Height decoding exceeded chance in all five sessions (37–73% balanced accuracy; chance = 33%). To test whether the spatial signal persists beyond sensory and kinematic confounds, we removed variance associated with measured visual, kinematic, and slow time-drift features from each unit’s firing rate and repeated the decoding. Residualized accuracy remained above chance across all sessions (38–57%), showing that the population spatial code is not reducible to these covariates. Finer-grained 9-zone decoding also exceeded chance in M2 for both raw and residualized activity (Figure 2E), though insufficient zone coverage in M1 precluded this analysis.

Spatial modulation maps for the highest-POS-encoding units showed localized hot spots within the arena (Figure 2G), while low-ranked units exhibited spatially uniform firing. Although visually striking, these maps may reflect a combination of visual, kinematic, and positional variables rather than a dedicated place code. The decoding and spatial mapping analyses indicate that mSTS population activity carries information about which vertical level the animal occupies and, in the better-sampled subject, about arena location within each level.

### Reference-frame encoding in mSTS favors allocentric over body-centric and head-centric coordinates

We next asked which coordinate frame best captured encoding of displacement and velocity variables during free movement. Describing movement relative to the world, body, or head requires a reference frame (RF): egocentric coordinates suffice for visuomotor control, but allocentric (world-fixed) coordinates are needed for tracking locations relative to stable environmental landmarks. We fit three reference-frame models – allocentric, body-centric, and head-centric – to 1,404 units across five sessions (Figure 3A; see Methods for model specification). Allocentric encoding was the best-fitting model for 50% of units, compared to 37% body-centric and 13% head-centric (Figure 3B). On the scatter of allocentric versus body-centric ΔR^2^, most strongly tuned units fell below the identity line (diagonal), confirming allocentric dominance at the single-unit level (Figure 3D). This pattern was observed in both multi-unit and single-unit activity, appeared in both electrode banks, and was most pronounced in the strongly tuned tail of the population (Figure 3C). At the unit level, ventral bank showed larger encoding magnitudes across all three frames (Mann–Whitney p < 0.001 for each frame; e.g., allocentric: ventral median ΔR^2^ = 0.059, dorsal = 0.026), whereas winner proportions did not differ detectably between banks (χ²(2) = 1.5, p = 0.47; Figures 3B,F and Extended Data Fig. 3); because units were nested within sessions and animals, these bank comparisons should be interpreted cautiously. These results indicate that mSTS activity is better explained by allocentric than by body- or head-centric displacement and velocity variables overall, with substantial body-centric contributions, consistent with mixed reference-frame coding during free behavior.

**Figure 3.**
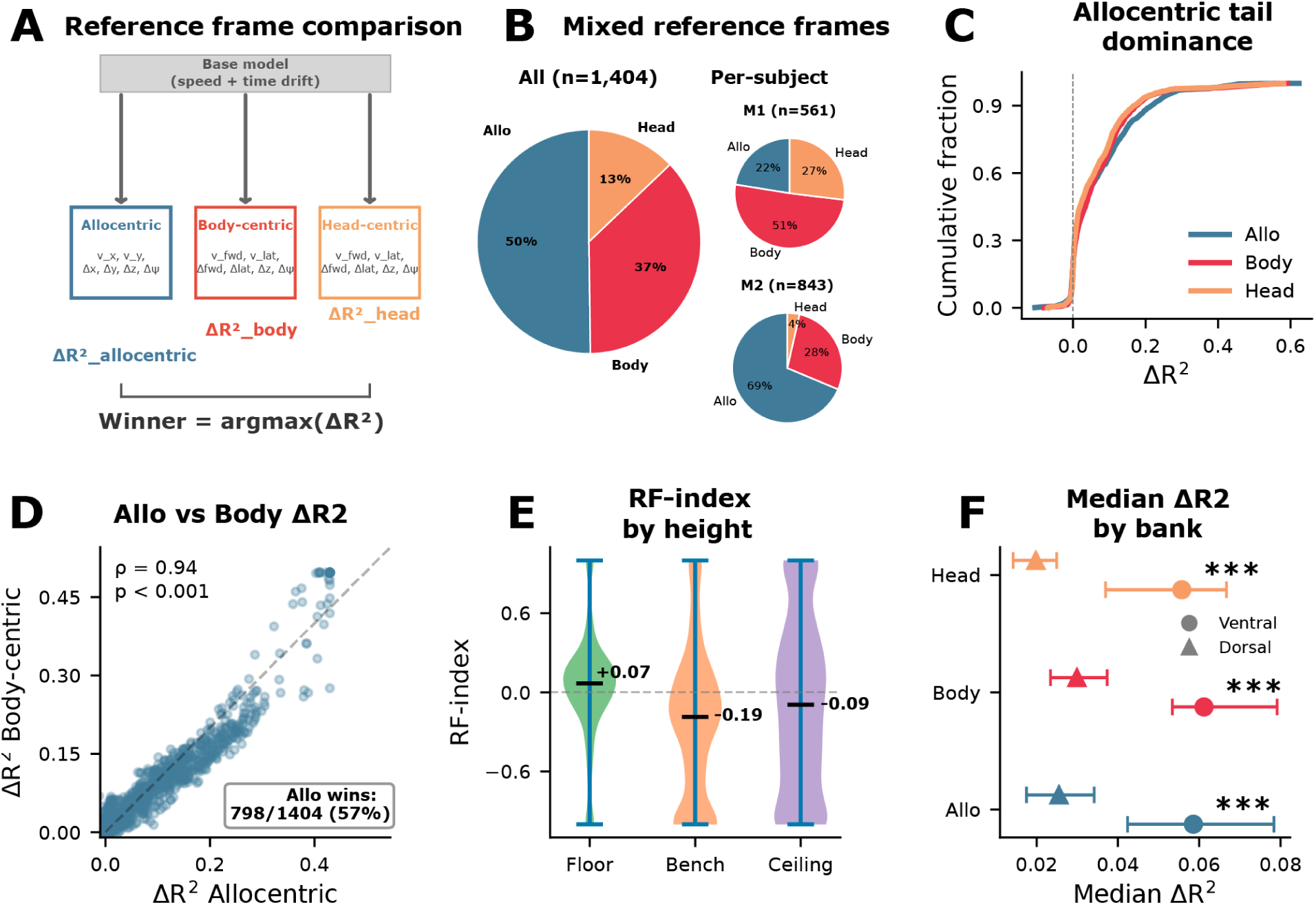
Reference-frame encoding in mSTS favors allocentric over body-centric and head-centric coordinates. **a.** Schematic of the reference-frame comparison. Three encoding models expressed velocity and displacement features in allocentric (world-fixed), body-centric, or head-centric coordinates (6 features each), sharing a common base model (speed + time drift). The winner frame for each unit was the frame with the largest ΔR^2^ above base. **b.** Winner-frame proportions as pie charts for pooled data (left) and by subject (right). Allocentric encoding was the most frequent winner overall (M2: 68.7% allo; M1: 50.6% body-centric), with both coordinate frames represented across subjects (n = 1,404 units, 5 sessions, 2 subjects). **c.** Cumulative distribution of ΔR^2^ by reference frame. Allocentric and body-centric curves show dominance in the upper tail. **d.** Allocentric ΔR^2^ (x-axis) versus body-centric ΔR^2^ (y-axis) for each unit. Most strongly tuned units fall below the diagonal, indicating allocentric dominance at the single-unit level. **e.** Height-dependent reference-frame index. Violin plots show the distribution of per-unit reference-frame (RF) index across height tiers (floor, bench, ceiling). Positive values indicate allocentric-dominant coding, negative values indicate body-centric-dominant coding, and values near 0 indicate mixed coding. Violin width indicates density, and horizontal bars mark medians. **f.** Median ΔR^2^ by reference frame and bank (dot = median, bars = 95% CI). Ventral bank showed significantly stronger encoding across all three frames (Mann–Whitney p < 0.001), but the relative ordering (allocentric > body-centric > head-centric) was preserved in both banks.

This overall allocentric preference was driven by M2, who showed strong allocentric dominance (68.7% allo, 27.8% body, 3.6% head). M1 showed a different balance, with body-centric coding predominating (22.5% allo, 50.6% body, 26.9% head). However, M1’s encoding magnitudes were lower; small differences near zero may reflect noise rather than a true frame preference. This between-subject difference may partly reflect differences in arena coverage, because M1 explored a smaller portion of the arena and therefore provided less allocentric spatial variance to the encoding model. Importantly, both reference frames were represented in both subjects (22.5% allocentric in M1; 27.8% body-centric in M2). The coexistence of both coordinate frames, together with subject-level shift, raises the question of whether frame preference is fixed or depends on behavioral context.

To test this, we asked whether reference-frame preference shifts with vertical context, since animals use different coordinate strategies at different heights (e.g., navigating open floor space versus climbing structured surfaces). We stratified the arena into three height tiers (floor, bench, ceiling) using fixed center-of-mass (COM)-height boundaries and re-fit encoding models independently within each tier (see Methods; Figure 3E). To move beyond binary winner classification, we computed a continuous RF-index contrasting allocentric and body-centric unique variance for each unit (see Methods). The RF-index shifted monotonically from allocentric-dominant on the floor (median RF-index = +0.60) through neutral at bench height (+0.02) to body-centric at the ceiling (-0.24) (Figure 3E). This context-dependent shift parallels a longstanding insight from primate positional behavior research. Differences between floor exploration and climbing may reflect changes in the relative importance of external landmarks versus posture- and body-centered control variables^41–43^. Together, these results indicate that reference-frame weighting for displacement and velocity variables is not fixed, but shifts with vertical context. While variance decomposition reveals what individual neurons encode, it does not address how the population jointly organizes these variables – a question relevant to what downstream circuits could read out.

### mSTS neurons encode discrete behavioral syllables with sustained temporal structure and structured population tuning

To test whether mSTS encodes discrete behavioral structure beyond continuous kinematics, we examined neural responses aligned to transitions between behavioral syllables (12 classes from unsupervised pose segmentation; see Methods). Onset-aligned population responses showed systematic modulation at syllable boundaries, with a wave of action across units sorted by peak latency (Figure 4A). We decoded syllable identity from population firing rates using multinomial logistic regression with temporal cross-validation (5-fold, 2 s embargo). Decoding exceeded chance in four of five sessions (19–25% balanced accuracy; chance = 10%; permutation p < 0.01), with one M1 session near chance (9.9%).

**Figure 4.**
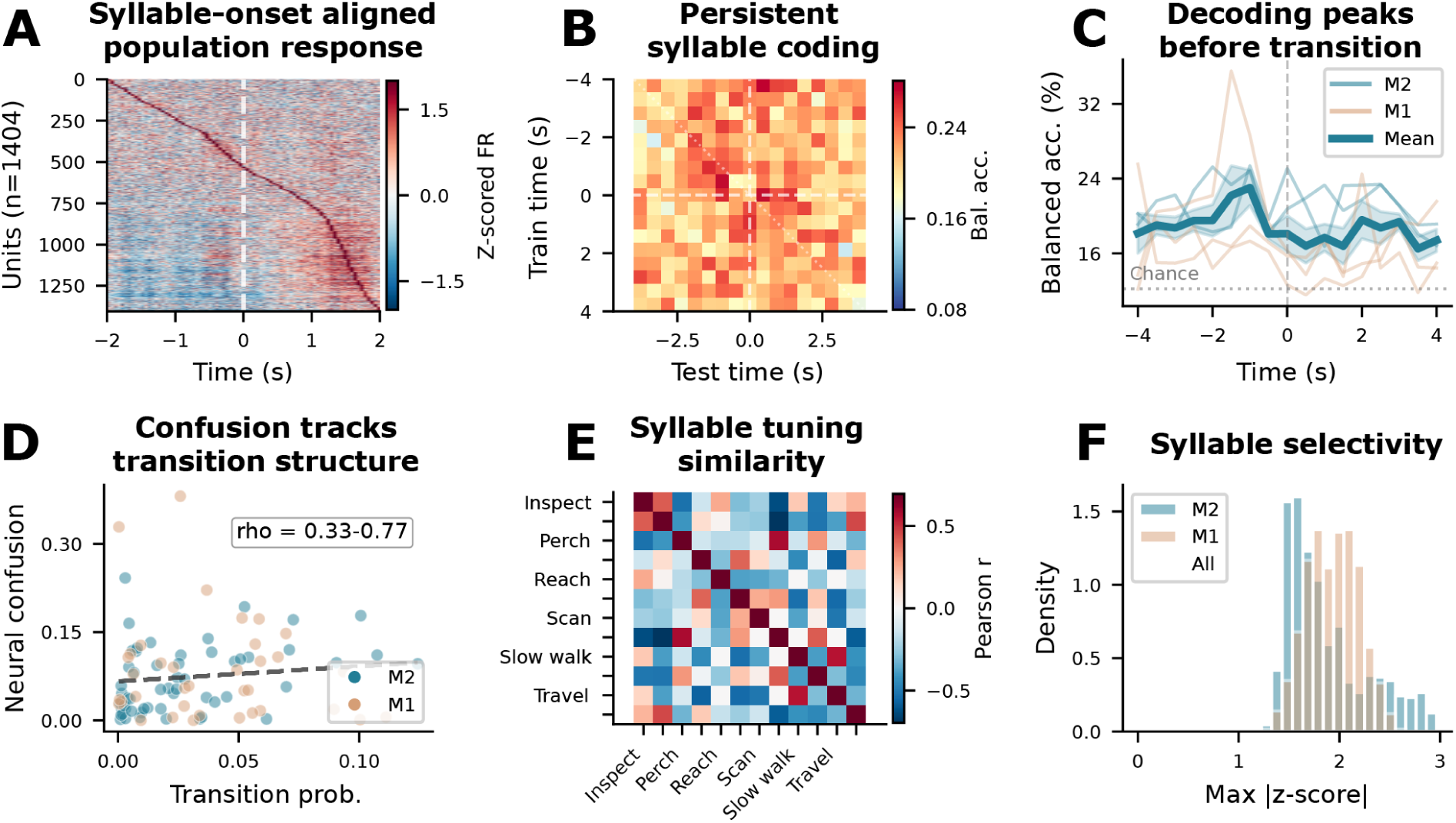
mSTS neurons encode discrete behavioral syllables with sustained temporal structure and structured population tuning. **a.** Onset-aligned peri-event time histogram (PETH). Rows = units (n = 1,404) sorted by peak latency; columns = time relative to syllable onset (±2 s, 50 ms bins). Z-scored firing rates reveal a temporal wave of population-level modulation at behavioral transitions. **b.** Temporal generalization matrix (TGM). Exemplar session (M2): a classifier trained to decode syllable identity at one peri-transition time point was tested at all other time points. Above-chance accuracy extends broadly off-diagonal, indicating a sustained or temporally generalizing syllable-related structure rather than a purely transient onset response. **c.** TGM diagonal accuracy (same-time decoding) for all five sessions (thin lines, colored by subject; bold = mean ± SEM). All sessions exceed chance (dashed). **d.** Neural confusion tracks behavioral transition structure. Each dot represents one off-diagonal syllable pair, with transition probability (x-axis) plotted against neural decoding confusion (y-axis), colored by session. Syllables that frequently follow each other in the behavioral stream are more often confused by the neural decoder (Mantel ρ = 0.3–0.8, permutation p < 0.05 in all sessions). **e.** Syllable tuning similarity. Pearson correlation of population z-scored tuning profiles across all 12 discovered syllables (pooled across sessions; decoding analyses in panels A–D use the 10 most frequent per session). Correlated blocks indicate syllable groups that recruit similar neural subpopulations; anti-correlated blocks indicate distinct populations. **f.** Per-unit selectivity (maximum |z-score| across syllables) distribution for each subject (M1, M2). Most units show moderate selectivity (peak ≈ 1.5 z), consistent with distributed mixed coding.

To distinguish persistent state coding from transient onset modulation, we asked whether a decoder trained at one moment around the transition generalized to other moments in time (temporal generalization matrix^44^, TGM). We trained classifiers at each time point and tested generalization across all other time points in a ±4 s peri-transition window. If syllable coding is transient, only classifiers trained and tested at the same lag should succeed (a thin diagonal); persistent coding predicts broad off-diagonal generalization. Across all five sessions, above-chance accuracy extended well beyond the diagonal (Figure 4B), with mean diagonal accuracy exceeding chance in every session (15–21% vs 10–14% chance; Figure 4C). This broad generalization is consistent with a sustained representation of the current syllable across the peri-transition window.

We next asked whether the neural code reflects the statistical structure of behavior. Neural confusion tracked behavioral transition adjacency across sessions (Mantel rho = 0.3–0.8, permutation p < 0.05; Figure 4D): syllables that frequently follow each other in behavior were more often confused by the neural decoder. Population tuning profiles showed structured similarity among syllable representations (Figure 4E) – stationary postures (Rest, Pivot, Scan) were encoded by correlated subpopulations, while active locomotion syllables recruited distinct populations. Per-unit selectivity was moderate (median peak |z| ≈ 1.5; Figure 4F), and syllable coding was present across both sulcal banks, with no consistent dorsoventral dissociation across subjects (Extended Data Fig. 4F, Extended Data Fig. 5A). These results are consistent with a distributed, sequence-informed code for discrete behavioral state.

### mSTS population activity jointly organizes spatial location and behavioral state with structured transition dynamics

We next asked whether spatial context and behavioral state share a common population code. We summarized population activity with supervised low-dimensional embeddings using CEBRA^45^ and compared key results against PCA baselines (see Methods for model families and cross-validation procedures). In the Spatial-lite embedding, trained without vertical-position labels, height varied smoothly along the low-dimensional population space (Figure 5B). In the Behavior embedding, which was trained on continuous spatial and kinematic variables rather than syllable labels, behavioral syllables nonetheless occupied partially distinct subregions (Figure 5C). Decoding from these low-dimensional embeddings yielded 54–78% height accuracy (chance = 33.3%; Figure 5D) and 17–27% syllable accuracy (chance = 9–14%; Figure 5E), with PCA baselines performing comparably. This indicates that both spatial and behavioral variables are recoverable from population activity and that the result does not depend on a specific nonlinear embedding method. Together, these embeddings indicate that spatial and behavioral structure are jointly recoverable from population activity.

**Figure 5.**
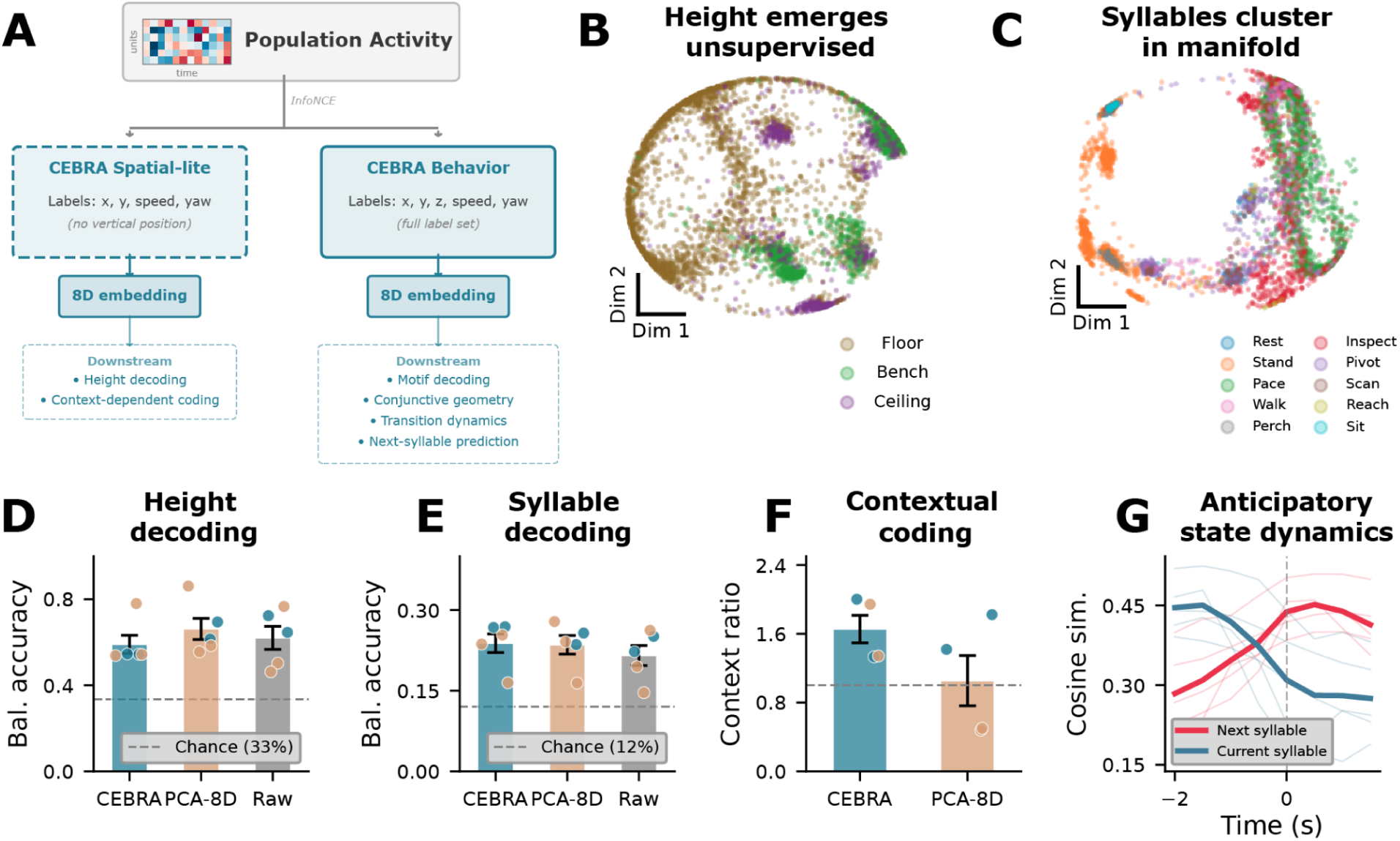
mSTS population activity jointly organizes spatial location and behavioral state with context-dependent coding and structured transition dynamics. **a.** Schematic of the CEBRA embedding analysis. Population firing rates (500 ms bins, z-scored) were embedded into 8D latent space using two separate CEBRA model families: a Spatial-lite model supervised with horizontal position and movement variables (com_x, com_y, speed, yaw rate; excluding vertical position) for height-related claims and context-dependent coding, and a Behavior model supervised with the full continuous label set, including vertical position (com_x, com_y, com_z, speed, yaw rate), used for downstream syllable-decoding and transition-dynamics analyses. All quantitative analyses used leave-one-fold-out temporal cross-validation with 2 s embargo, with CEBRA models fit independently on each training fold. **b.** CEBRA-Spatial-lite 2D projection of M2 population activity (443 units, session 1), colored by height tier (floor/bench/ceiling). Despite the exclusion of vertical position from training labels, height structure is organized along the embedding. **c.** CEBRA-Behavior 2D projection colored by behavioral syllable identity (10 classes). Syllables occupy distinct subregions, indicating joint spatial-behavioral organization. **d.** Height decoding (3 classes: floor, bench, ceiling). Bars show cross-session mean balanced accuracy (± SEM) for CEBRA-Spatial-lite 8D kNN, PCA-8D kNN, and raw logistic regression; dots show individual sessions (n = 5, colored by subject). All methods exceed chance (33.3%, dashed). Height: CEBRA-SL 54–78%, PCA-8D 56–86%, Raw 46–72%. **e.** Syllable decoding. Same format as (D). CEBRA-Behavior 8D kNN, PCA-8D kNN, and raw logistic regression. CEBRA-B 17–27%, PCA-8D 16–28%, Raw 15–26% (chance = 9–14%). **f.** Context-dependent coding. Between-context centroid distances (same syllable at different height tiers) versus within-context distances in Spatial-lite embedding space (trained without height labels), summarized as a context ratio. Ratios 1.33–3.73 across all five sessions (permutation p < 0.001), confirming that the same behavior occupies different population space at different locations. PCA-8D showed the same pattern in M2 (ratios 1.42, 1.82). All 8 CEBRA dimensions encoded both syllable and height simultaneously (η² > 0.02 for both variables in all sessions), consistent with a joint state-space representation. **g.** Anticipatory drift. Cosine similarity between the neural state and the next-syllable centroid (red) versus the current-syllable centroid (blue) in a ±2 s window around behavioral transitions. The pre-transition crossover indicates structured anticipatory dynamics (pre-ramp p < 0.01 in 4/5 sessions).

If space and behavior are jointly organized, the same syllable performed in different spatial contexts should occupy different regions of the low-dimensional population space. We tested this by comparing how far apart the average population patterns were when the same syllable occurred at different height tiers (i.e., using centroid distances; Figure 5F). Between-context distances significantly exceeded within-context spread across all five sessions (ratios 1.3–3.7; permutation p < 0.001), indicating that the population representation of a behavior shifts with the animal’s vertical position. The same context-dependent organization was observed in PCA space for M2, indicating that the effect did not depend on a specific nonlinear embedding method. We further assessed whether this joint coding was confined to specialized dimensions by computing the variance explained (eta-squared) by each variable per embedding dimension. All eight dimensions carried variance related to both syllable identity and height in every session, indicating distributed low-dimensional structure for both variables rather than an obvious separation across dimensions (Figure 5F). These results are consistent with a joint low-dimensional representation of behavioral and spatial variables, such that similar behaviors are expressed differently across spatial contexts.

To assess temporal organization around behavioral transitions, we computed the cosine similarity between the neural state and the upcoming syllable centroid in a ±2 s window around transitions (Figure 5G). The neural trajectory drifted toward the upcoming syllable centroid before the behavioral transition (permutation p < 0.01 in 4/5 sessions), consistent across M2’s two sessions and detectable at reduced magnitude in M1. Neural state velocity was also elevated at transitions (42–117% above baseline; Extended Data Fig. 5D). The neural population thus shows structured temporal dynamics around behavioral transitions, with activity patterns changing before overt state changes.

## Discussion

Here we characterized mSTS neural activity during unconstrained behavior using wireless depth recordings, markerless 3D pose tracking, and unsupervised behavioral segmentation in freely moving macaques. We made five principal discoveries. First, static 3D position in the arena explained more unique variance in mSTS firing rates than any single measured visual or kinematic feature group. Second, in separate reference-frame models based on displacement and velocity variables, allocentric coordinates outperformed body- and head-centric coordinates overall. Third, reference-frame preference was flexible across the 3D environment, shifting from allocentric-dominant near the floor toward more body-centric coding during vertical exploration. Fourth, mSTS populations carried information about discrete behavioral syllables with sustained, sequence-related structure across time. Fifth, population activity showed joint organization of spatial context and behavioral state, such that the same behavior occupied different neural-state regions at different locations, and these dynamics were temporally organized around behavioral transitions. Together, these findings suggest that mSTS activity carries joint information about position-related variables, reference-frame structure, and behavioral state during natural behavior.

The central population-level finding is that the same behavioral syllable occupies different regions of the neural manifold depending on spatial context (Figure 5F). This organization has a clear computational consequence. Behaviorally similar actions can have different sensory and motor consequences depending on where they occur. A code that represented behavior without context would lose that distinction (e.g., a reach performed on the ground requires a different postural correction than one performed while suspended), whereas the joint organization we observe could provide downstream circuits with context-specific information about ongoing state. Coding was distributed across both sulcal banks without dorsoventral segregation, consistent with mixed selectivity as a general principle^46^ rather than modular organization.

The anticipatory drift we observe – in which the neural trajectory shifts toward the upcoming syllable’s centroid before the overt behavioral transition – suggests that mSTS has access to prospective information about imminent state changes. One candidate mechanism is efference copy from premotor or prefrontal circuits known to encode hierarchical action plans^9,12^, which could bias mSTS population dynamics ahead of movement execution. Alternatively, recurrent dynamics within STS itself, possibly shaped by the sequential statistics of the animal’s own behavioral repertoire, could generate predictive drift without requiring an external motor signal. Disambiguating these accounts will require simultaneous multi-region recordings or targeted perturbation during free behavior.

These findings are also consistent with the idea that brain regions linked to social cognition may be recruited because they perform computations that are useful in many settings, not only social ones^35^. The signals we observe in solitary mSTS – arena position, flexible reference-frame weighting, sustained behavioral-state coding, and a joint population organization of action-in-context – are the kinds of variables that could support agent monitoring in a shared physical environment. In that sense, the present data do not demonstrate that mSTS is exclusively social; rather, the computations observed here during solitary behavior are consistent with a domain-general system that could also serve social functions.

Several limitations constrain interpretation. First, the dataset comprised 1,404 units recorded across five sessions in two animals, so many statistical comparisons were performed at the unit level within a hierarchically nested design (units within sessions within animals). We therefore treat these analyses as evidence for robust structure within the recorded dataset, not as strong animal-level inference. Second, M2 showed strong effects across analyses, whereas M1 showed weaker signals reflecting restricted arena coverage; M1 therefore provides compatible but limited replication rather than independent confirmation. Our visual feature set captures geometric variables such as gaze direction and looming-like motion, but not full-field optic flow or scene texture; a richer visual model could reduce the apparent dominance of position, although the residualized spatial signal provides a lower bound that is independent of measured visual and kinematic covariates. Further, free behavior couples visual, proprioceptive, and vestibular inputs, so variance decomposition can separate unique contributions but cannot establish causality. Finally, the supervised CEBRA embeddings are trained on continuous spatial and kinematic labels, so the learned manifold organization partly reflects the training variables; however, context-dependent coding was confirmed in unsupervised PCA space, and a self-supervised CEBRA-Time variant recovered partial spatial and behavioral structure without any behavioral labels (Extended Data Fig. 5B).

Our findings highlight the central promise of primate neuroethology^3^. Studying the brain during natural behavior reveals computational patterns that constrained paradigms systematically miss^6,12^. mSTS has been defined by its visual responses to faces, bodies, observed actions, and even coding of reward and social prediction^21,24^, yet when the animal is free to move, position-related variables and behavioral state account for a substantial portion of its measured encoding structure. A natural next step is to test whether social interaction recruits or reshapes this joint organization of spatial context and behavioral state so that it tracks a partner alongside the self. Whatever its full role, our understanding of mSTS should evolve from a purely observer-focused hub for social perception^19,24,25,47–50^ to include a region whose activity carries information about where the animal is and what it is doing – signals that could support real-time coordination with others during natural behavior.

## Methods

### Animal Subjects

Two adult rhesus macaques (M. mulatta; one male and one female) participated in the experiment. Neural activity was recorded from each animal individually. Animals were trained using positive reinforcement by clicker training to enter a primate chair (Hybex Innovations), which was used briefly to stabilize the head (up to 5 min) while positioning the neural data logger. Animals were trained to enter and exit the open-field tracking arena where all experiments took place. Experiments took place between 10:00 AM and 1:00 PM from May to July 2024 across 30 experimental days, with ample rest periods between sessions. The arena also served as an enrichment space; foraging materials (seeds, nuts) were scattered throughout to encourage naturalistic exploration and multi-level space use. All procedures were approved by the University of Pennsylvania Institutional Animal Care and Use Committee (IACUC).

### Wireless Neural Recording with Multi-Electrode Sulcal Depth Arrays

We recorded extracellular neural activity from the right middle superior temporal sulcus (mSTS) of two rhesus macaques (M1 and M2) using 128-channel NeuroNexus depth electrode arrays (M4x8-6mm-550-500-177; 4 shanks x 8 contacts per probe, 550 µm inter-contact spacing, 500 µm inter-shank spacing, 177 µm^2^ contact area). Two probes were implanted per bank (A1/A2 = ventral, B1/B2 = dorsal), with 800 µm anterior–posterior spacing between probe platforms (A2/B2 anterior, A1/B1 posterior), yielding 128 electrode sites total (64 ventral bank, 64 dorsal bank). Neural signals were transmitted wirelessly via skull-mounted data loggers (Deuteron Technologies) while animals freely navigated a hexagonal arena (height 2.4 m). Sessions lasted 30–60 minutes across five recording sessions (3 M1, 2 M2).

### Neural Preprocessing and Semi-Automated Spike Sorting

Spike sorting followed a four-stage semi-automated pipeline. Raw extracellular signals were bandpass-filtered (250 Hz high-pass, 4th-order Butterworth) and denoised using median-based common average referencing (CAR) applied separately within each electrode bank. Channels with confirmed physical damage (M1: 24 channels on broken shanks identified during implantation) or chronic noise (flagged via automated waveform quality analysis across sessions; Extended Data Fig. 1) were excluded from the CAR reference signal.

Spikes were detected at ∼3.7 median absolute deviations (MAD) below zero with a 1.25 ms refractory dead time, and waveforms (32 samples at 32 kHz, 8 pre-threshold) were extracted. Detected spikes were then sorted using the scan valley-seeking algorithm in 3D PCA space (first three principal components by variance explained; Plexon Offline Sorter V4), with Parzen radius scanned from 0.4 to 2.0 and cluster separation optimized via the J3 statistic. A trained researcher reviewed all clusters, merging or splitting as needed based on waveform morphology, ISI distributions, and autocorrelation structure.

Sorted clusters that survived manual quality check were post-processed automatically: clusters with ≥5,000 spikes were retained as single-unit (SUA) candidates, a minimum count ensuring stable waveform template estimation. Smaller clusters were merged into the nearest SUA template if the normalized RMS waveform distance was <0.25, or otherwise assigned to multi-unit activity (MUA). MUA for each channel comprised all unsorted spikes, orphan clusters, and spikes failing template rescue (distance ≥0.3). ISI violations (<1 ms) were flagged for units exceeding a 2% threshold; units exceeding this threshold were reclassified as MUA.

### Open-field Tracking Arena: Multi-camera Pose Estimation and Coordinate Frame Calibration

The arena was monitored by 30 WhiteMatter e3Vision cameras with hardware-triggered acquisition at 40 Hz, except for one 30 Hz session (hooke_0609). The number of usable cameras varied by session, typically from 26 to 30, depending on calibration quality and camera dropout. Camera intrinsics and extrinsics were estimated from checkerboard calibration using multical (https://github.com/oliver-batchelor/multical).

Two-dimensional keypoints were detected in each camera view using RTMDet-L for detection and RTMPose-L for pose estimation, yielding 17-keypoint skeleton detections per frame per camera. Three-dimensional body pose was reconstructed with batch direct linear transform and RANSAC-based outlier rejection. The reconstruction required at least six camera views per joint, keypoint confidence at least 0.3, inter-ray angle at least 3 degrees, and reprojection error at most 5 pixels for inlier classification. Joints falling outside the convex hull of camera centers were rejected. Median reprojection error across sessions was 0.35 pixels (Extended Data Fig. 1).

Reconstructed 3D coordinates were smoothed with a median filter (0.2 s window) followed by a mean filter (0.4 s window). Anatomical constraints enforced bone-length consistency (rejection at 3x MAD from the per-edge median length) and hinge-joint angle limits (15°–170° for elbows and knees only; remaining joints unconstrained).

A rotation matrix R_total_ was computed to align the coordinate frame such that the vertical axis corresponded to the direction of gravitational acceleration (estimated from the floor plane normal). All pose data and camera extrinsic parameters were rotated by R_total_ to ensure consistent alignment between animal trajectories and arena geometry. Arena center and wall geometry were estimated from the rotated camera positions by fitting planes to the hexagonal boundary; these defined the allocentric reference frame used in subsequent analyses.

### Neural and Video Data Synchronization

Neural logger clocks were synchronized to video using LED pulse alignment. An LED visible to the camera system was pulsed at session start and end; 5–8 pulses spaced across each session provided a broad temporal baseline for drift estimation. For each logger, a direct affine mapping (logger seconds to video frames) was estimated by regressing logger-side TTL edge timestamps against manually marked video LED onset times, with the first detected light pulse serving as the temporal origin. LED identity was verified by minimizing residuals across early and late pulse clusters. This procedure achieved sub-frame alignment (typically 0.2–0.6 frames, ∼5-11 ms at 40 Hz) over recordings lasting up to two hours (Extended Data Fig. 1).

### Automated Behavioral Syllable Segmentation

Discrete behavioral syllables were identified from 3D pose dynamics using an unsupervised segmentation pipeline adapted from [^39^] to a 17-keypoint primate skeleton. To control for body-size differences between animals, 3D keypoint coordinates were normalized per animal using the geometric mean of median torso height and median shoulder width (97.5th percentile scalar). All-to-all inter-keypoint Euclidean distances were computed for 12 body joints (face joints excluded), yielding 66 distance features per frame. These were reduced via PCA to 20 components. To ensure syllables captured behavior rather than individual morphology, the point-biserial correlation of each PC with binary animal identity (M2 vs. M1) was computed, and PCs with |r| > 0.3 were removed (a conservative threshold chosen to err toward excluding identity-contaminated components while retaining sufficient behavioral variance). This left 6 PCs that were uncorrelated with animal identity, consistent with the low-dimensional behavioral representations used in [^39^].

A Morlet continuous wavelet transform was applied to each retained PC at 20 geometrically-spaced frequencies (0.5–12.0 Hz; amplitude output, per-frequency z-normalization), producing 120 wavelet features. Per-joint heights (signed distance to floor plane; 13 joints, scale factor x 3.0 relative to z-scored wavelet features) and per-joint speeds (3D velocity magnitude; 13 joints, scale factor x 2.0) were concatenated, yielding 146 total features per frame. Heights were upweighted to capture vertical stratification (floor vs. bench vs. ceiling); speeds were upweighted to distinguish stationary from dynamic behaviors (Extended Data Fig. 4).

Features were embedded into 2D space using UMAP (n_neighbors_ = 15, min_dist = 0.1, cosine metric), fit on 47,722 training frames sampled with coverage-based balancing across sessions and animals (maximum 40% of frames from any single animal). Remaining frames were embedded into this reference space to obtain syllable labels across the full dataset. The 2D embedding was rasterized onto a 300 x 300 grid, Gaussian-smoothed (σ = 1.5), and segmented via watershed on the negative density landscape with auto-tuned peak separation. This yielded 26 initial density peaks, which were consolidated to 12 final syllables via hierarchical merging (Ward linkage on trajectory embeddings, Euclidean distance), guided by inspection of merged clusters to ensure systematically distinct behavioral categories. A median filter (40 frames = 1 s at 40 Hz) was applied to the ethogram, and bouts shorter than 1 s were absorbed into neighboring syllables, yielding 12 behavioral syllables (median bout duration 1.50 s, IQR 0.43–3.40 s).

For neural analyses, the 10 most frequent syllables in each session were retained; rarer syllables were collapsed to an “other” category and then excluded, yielding a 10-class classification problem. Syllable labels were aligned to 500 ms neural bins by assigning each bin the nearest behavioral frame within 500 ms of the bin center. Bins without a behavioral frame in this window were marked missing and excluded.

### Shared neural encoding framework

Neural activity was modeled as a linear function of behavioral features using ridge regression. For each unit, firing rate was predicted from the z-scored design matrix, with regularization strength chosen on training data only.

Model performance was evaluated with blocked temporal cross-validation to account for autocorrelation in neural time series. Data were divided into contiguous 30 s blocks assigned cyclically to folds, and a 2 s embargo was excluded from training on either side of each test block. Regularization strength was selected by nested cross-validation on a grid of 20 log-spaced alpha values from 10^-2^ to 10^4^. Features were z-scored within each fold using training-set statistics, and cross-validated R2 was computed from pooled held-out predictions. Neurons were included if they had at least 150 valid bins after lag trimming and exclusion of missing values.

### Variance decomposition and spatial decoding

Behavioral features were organized into four groups. VIS contained five geometric visual proxies: three components of head-forward gaze direction, distance to the nearest wall along the horizontal gaze direction, and a looming-like visual motion term. BODY_KIN contained forward velocity, lateral velocity, and 3D speed. HEAD_KIN contained yaw, pitch, and roll velocity, plus head-body yaw offset. POS contained center-of-mass x, y, and z coordinates in the rotated arena frame. VIS, BODY_KIN, and HEAD_KIN features were expanded with temporal lags spanning ±1.0 s at 500 ms resolution; POS was entered as an unlagged linear regressor. A linear time-drift covariate was appended. Figure 2 used 10-fold cross-validation.

To quantify the unique contribution of each feature group, we used the regressor-shuffle approach of Testard et al. (2024). For each neuron, alpha was selected once on the full model and then held fixed for all follow-up fits to ensure that differences in ΔR² reflect feature-group contributions rather than changes in regularization. One feature group at a time was block-permuted in 10 s chunks to destroy its alignment with the neural response while preserving local autocorrelation. The unique contribution was the average decrease in cross-validated R2 relative to the intact model. Empirical one-sided p-values were computed from 100 shuffle iterations and corrected across neurons with the Benjamini-Hochberg procedure. A feature group was considered significant for a neuron if the FDR-corrected p-value was below 0.05, unique ΔR² was at least 0.005, and full-model R2 exceeded 0.01.

To test whether the population carried sufficient spatial information to decode location, we trained multinomial logistic regression models on the full population firing-rate vector. The arena was discretized into three height tiers (floor, bench, ceiling) defined by tercile boundaries of floor-relative center-of-mass height (33.3rd and 66.7th percentiles of com_z pooled across all sessions). For finer-grained decoding, each height tier was further divided into three 120-degree allocentric angular sectors (computed from arctan2 of horizontal center-of-mass position relative to the arena center), yielding nine composite zones. Decoding used 5-fold blocked temporal cross-validation with the same structure as the encoding models. Folds were valid only if both train and test sets contained at least one example of each class. To test whether spatial information persisted after accounting for covariates, we residualized each unit’s firing rate against VIS, BODY_KIN, HEAD_KIN, and time-drift regressors, then repeated the decoding. Observed balanced accuracy was compared with a null distribution from 100 block-permutations of spatial labels.

### Reference frame analyses

Three encoding models expressed velocity and displacement features in allocentric, body-centric, or head-centric coordinates (6 features each), sharing a common base model (speed plus time drift). Each frame-specific model selected its own regularization parameter by nested cross-validation. Because all three frame models had identical dimensionality, differences in cross-validated R2 above base reflect coordinate-frame preference rather than model complexity. The winner frame for each neuron was the model with the largest improvement over base. Figure 3 used 5-fold cross-validation.

To test whether frame preference changed with vertical context, we repeated the reference-frame comparison independently within each height tier. Temporal folds were constructed on the full session timeline and intersected with bins belonging to each tier, so that temporal blocking was preserved while train and test sets were restricted to one height level. Tiers with fewer than 50 bins per unit were excluded, and a unit-tier estimate was considered interpretable when at least two valid folds remained, at least 20 test bins remained in the tier, and the firing-rate variance exceeded a floor of 1e-6. The paired tier analysis was restricted to the 203 units interpretable across all three tiers. Floor-to-ceiling RF-index differences were assessed with a paired Wilcoxon signed-rank test.

### Syllable decoding and population tuning

Population activity was decoded into syllable identity with multinomial logistic regression using balanced class weights. Each 500 ms neural bin was treated as one sample. Decoding used 5-fold blocked temporal cross-validation with contiguous time blocks and a 2 s embargo on either side of the test block. Features were z-scored within each fold using training-set statistics. Balanced accuracy was used to account for class imbalance. Statistical significance was assessed by comparing observed accuracy to a null distribution from 1,000 label permutations preserving the same fold structure. To assess temporal alignment, decoding was repeated at lags from -2.0 to +2.0 s.

To distinguish persistent state coding from transient onset responses, we constructed a temporal generalization matrix (TGM). For each syllable transition, we extracted a peri-event window spanning 4 s before to 4 s after onset. Classifiers trained at one time point were tested at all others using 5-fold stratified cross-validation over transitions with a 2 s embargo. Neural features were z-scored within each fold and reduced to at most 50 principal components when needed. Chance was defined as 1/n_classes. Diagonal accuracy was highest in the pre-transition window, consistent with anticipatory encoding of the upcoming syllable.

Neural confusion was compared with behavioral transition structure by computing the Spearman rank correlation between the symmetrized decoding confusion matrix and the empirical syllable transition probability matrix, with significance assessed by label-permutation test. Population-level tuning similarity was quantified as the Pearson correlation between z-scored tuning vectors across units. Per-unit selectivity was defined as the maximum absolute z-score across syllables. For onset-aligned visualizations, population firing rates were aligned to syllable onsets and z-scored per unit for display.

### Population manifold analysis

To characterize population-level organization, we applied CEBRA, a contrastive learning framework for nonlinear embedding of neural population activity. The input for each session was the z-scored firing-rate matrix across 500 ms bins and all recorded units. Two supervised model families were trained to avoid circularity. The Spatial-lite model was trained on continuous labels for horizontal center-of-mass position, locomotor speed, and head yaw rate (com_x, com_y, speed, yaw_rate) and was used for height-related claims and context-dependent coding because vertical position was not supplied during training. The Behavior model was trained on the full continuous label set including vertical position (com_x, com_y, com_z, speed, yaw_rate) and was used for syllable decoding and transition analyses. Both models used the offset10-model architecture, batch size 512, and 5,000 training iterations per session. For context-dependent coding, we used the 3D Spatial-lite embedding because this space was also used for visualization and preserved the effect robustly; PCA controls were computed in 8D to match the standard quantitative baseline. For quantitative analyses, each session was split into five contiguous temporal folds with a 2 s embargo, and a separate CEBRA model was fit on each training fold. Visualization embeddings were fit on all data and used for illustration only. Discrete labels were decoded from 8D CEBRA embeddings with k-nearest-neighbor classification (k = 5) and compared against PCA-8D and raw logistic regression baselines.

To test context-dependent coding, we computed centroids in 3D Spatial-lite embedding space for each syllable-by-height-tier combination (requiring at least 20 bins per cell) and compared between-context centroid distance with within-context spread. The context ratio was tested against a null generated by permuting tier assignments within each syllable (500 permutations). The nonlinear analysis used 3D visualization-quality embeddings; the PCA control used 8D to match the quantitative CEBRA decoding dimensionality.

To characterize transition dynamics, we extracted neural trajectories in 3D Behavior embedding space in a ±2 s window around each syllable onset, defined as the first bin of a new syllable following a different syllable. Syllable centroids were computed from all bins assigned to each syllable across the full session. We quantified whether the trajectory moved closer to the upcoming syllable centroid before transition onset using cosine similarity. Transition-aligned neural velocity was computed separately as the Euclidean distance between consecutive embedding states and compared with non-transition baseline bins after excluding all bins within ±2 s of any transition. Statistical significance for both anticipatory drift and velocity elevation was assessed with circular-shift null tests (500 shifts) preserving temporal autocorrelation. To test whether syllable identity and height tier were represented jointly or in separate latent dimensions, we computed eta-squared for each of the eight CEBRA dimensions separately for syllable identity and height tier. A joint state-space representation was inferred when most dimensions carried both variables.

### Statistical tests

Differences in ΔR² between dorsal and ventral banks were assessed with two-sided Mann-Whitney U tests. Differences in winner-frame proportions across banks were assessed with chi-squared tests of independence. Recording epochs were trimmed to manually annotated valid-inference windows before all analyses.

### Pseudoreplication controls

Multiple single units were recorded per electrode channel. To check that within-channel correlations did not inflate effect sizes, we repeated key analyses on one-per-channel subsets that retained the best single unit per channel per session. Results were also stratified by unit type where relevant.

### Software Availability

Analysis code can be found at: https://github.com/felipe-parodi/solo_mSTS_parodi2026. The encoding and decoding pipeline was implemented in Python 3.10 using scikit-learn, NumPy, SciPy, pandas, and matplotlib. Population manifold analysis used CEBRA v0.4.0 with PyTorch 2.1. Two-dimensional pose estimation used mmdetection and mmpose. Camera calibration used multical. Behavioral segmentation used PyWavelets, UMAP, and scikit-image. Spike sorting used Plexon Offline Sorter V4.

## Acknowledgements

We would like to thank the labs of Michael L. Platt and Konrad P. Kording for their valuable advice and support. This research was supported by NIMH, NIH, and NIA (R01MH095894, R01MH108627, R37MH109728, R21AG073958, R01MH118203, R56MH122819 and R01NS123054 to M.L.P).

